# Metabolic fingerprinting of 17 Brassicaceae species across three tissues

**DOI:** 10.64898/2026.04.17.719198

**Authors:** Felicia C. Wolters, Tina Woldu, Eric M. Schranz, Marnix H. Medema, Klaas Bouwmeester, Justin J. J. van der Hooft

**Author notes:** Corresponding author(s): Felicia Wolters, Marnix Medema & Justin van der Hooft. Shared last authors (Klaas Bouwmeester, Justin J. J. van der Hooft).

## Abstract

Plants produce the most diverse blends of specialized metabolites on earth. Natural products derived from plants are valuable resources for drug development, food chemistry, and crop resistance breeding. Phenotypes of specialized metabolite profiles can be captured by untargeted mass-spectrometry across species phylogeny, tissues, and genotypes. Here, we collected metabolic fingerprints of 17 Brassicaceae species across three tissues (paired leaf and root; flower) using liquid chromatography-tandem mass spectrometry (LC-MS/MS) in positive and negative ionization mode. Corresponding metadata has been refined for reuse according to ReDU guidelines, and for integration with public genomic and transcriptomic data. Standardization of *in vitro* growth conditions, and data processing workflows enables integration of acquired raw and processed data across platforms for single- and multi-omics analysis. Further, the inclusion of tissue-specific metabolic profiles across ploidy levels, as well as across crop species and wild relatives, makes this dataset a valuable resource for natural product discovery.

## Background & Summary

Plant specialized metabolite diversity has been studied extensively for natural product discovery with applications in human medicine, agrochemistry, and targeted plant breeding^1–3^. In plants, evolution of specialized metabolite biosynthesis is especially complex in polyploid species ^4,5^. Metabolic profiling revealed diversification of specialized compounds on the phylogenetic levels of plant families, clades, lineages, and species^3,6–8^. The Brassicaceae family comprises multiple economically important polyploid crops^4,5^, the model species *Arabidopsis thaliana*, and several crop wild relatives covered by available genome data resources in public repositories. The evolution of genome rearrangements following polyploidization and their effect on specialized metabolic pathway evolution have been studied extensively in this plant family^4,5^. In particular, glucosinolate biosynthesis exemplifies the evolution of a specialized compound class involved in plant resistance to biotic stress specific to the Brassicaceae^9–11^.

Deciphering complex biosynthetic pathways requires the acquisition and interpretation of molecular omics datasets, including untargeted metabolomics data. Integrating multiple omics data types is becoming an increasingly powerful strategy for discovering novel plant biosynthetic pathways^1^. However, the extensive diversity of natural product biosynthesis across species, genotypes, tissues, and environmental conditions poses significant challenges for integration of single or multi-omics data acquired in independent research.

Here, we present a paired tissue-specific metabolic fingerprinting dataset for 17 species in the Brassicaceae family across three tissues (leaf, root, flower), consisting of untargeted Liquid Chromatography-Mass Spectrometry (LC-MS/MS) data in positive and negative ion detection mode. This untargeted metabolomics dataset was acquired from paired leaf and root tissue of plants growing *in vitro* under standardized conditions in a climate chamber, and from flower tissue of plants growing in soil under greenhouse conditions. The presented dataset covers a range of crop species and wild relatives with publicly available sequenced genomes across three lineages in the Brassicaceae family (Brassicodae, Camelinodae, Arabodae)^5,12^ and the outgroup species *Aethionema arabicum* (Brassicales). Integration with public datasets is facilitated by a standardized experimental design and metadata format, sampling scheme, and modular downstream processing workflows. Raw and pre-processed data is provided in combination with extracted feature lists and annotations in the MetaboLights^16^ and MassIVE^17^ metabolomics data repository, and downstream processing workflows on GitHub and Zenodo.

## Methods

### Plant material and growth conditions

Seeds of plant genotypes listed in Table 1 were vapor-sterilized using chlorine gas in a desiccator for 3h, cold stratified at 8°C for seven days, and sown on half-strength Murashige and Skoog (MS) agar. Plants were grown for 25 to 35 days in a climate chamber (16:8h photoperiod, 140-180 µmol m2 s-1, 22 ± 2°C) until seedlings developed four mature leaves (PO:0007115). Plant positions in the climate chamber were randomized.

**Table 1.**
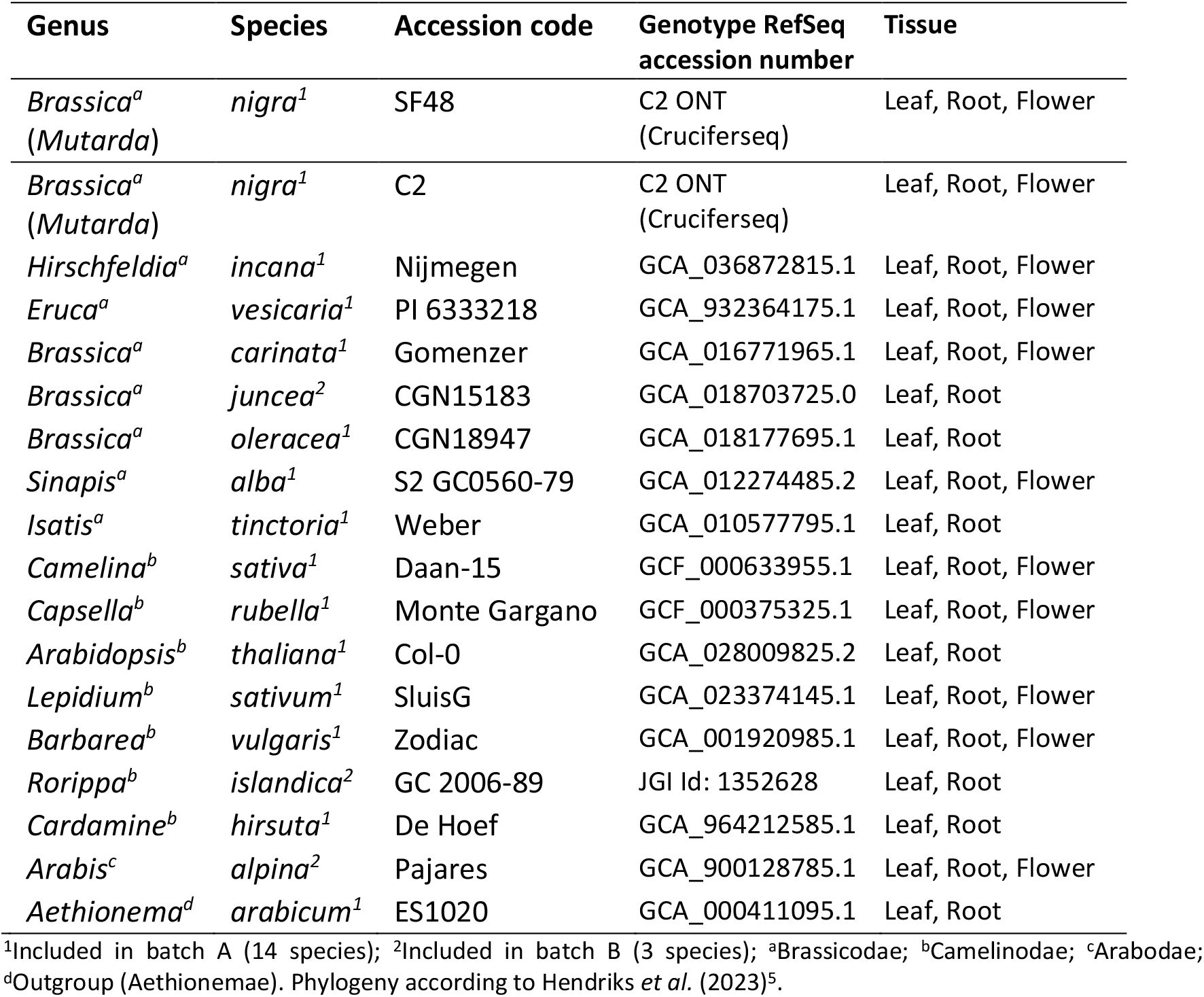
Species and genotypes included in LC-MS/MS dataset with reference to public genome assemblies on NCBI, JGI, and Cruciferseq, taxonomic order, and sampled tissue.

**Table 2.** Metabolite features detected across 14 species and three tissues in positive and negative ionization mode with consensus NP.Classifier annotations across at least two out of three annotation tools (SIRIUS, MS2Query, DreaMS)

| Tissue<sup>1</sup> | Positive ionization mode |  |  | Negative ionization mode |  |  |
| --- | --- | --- | --- | --- | --- | --- |
|  | Leaf | Root | Flower<sup>2</sup> | Leaf | Root | Flower<sup>2</sup> |
| Aligned feature list (unfiltered) | 5,106 | 2,912 | 4,222 | 4,278 | 2,155 | 3,164 |
| Consensus NPC.Pathway | 3,443 | 1,716 | 2,884 | 2,780 | 2,229 | 2,165 |
| Consensus NPC.Superclass | 2,045 | 920 | 1,758 | 1,638 | 706 | 1,327 |
| Consensus NPC.Class | 1,316 | 602 | 1,215 | 1,200 | 481 | 999 |

<sup>1</sup>Values given for total number of features detected in leaf and root tissue, including tissue-specific metabolite features, and features shared between leaf and root tissue.
<sup>2</sup>Raw LC-MS/MS data obtained from flower tissue samples was aligned and processed in separate batches from leaf and root tissue.

### Plant tissue sampling

Plants were harvested from agar pots, and leaf and root tissue were carefully separated, using a scalpel, excluding stem and cotyledons. Residual growth medium was removed from roots by washing with demineralized water. Harvesting was performed after 8-10 h of light exposure (between 1 and 3 PM), and collected tissues were immediately frozen in liquid nitrogen. Samples from three biological replicates per species consisting of 5-20 plants per replicate grown in independent tissue culture units were harvested and homogenized under liquid nitrogen using a mortar and pestle and stored at −80°C until further analysis.

### Metabolite extraction and LC-MS/MS analysis

Tissue samples were processed for metabolite extraction and analysis according to De Vos et al. (2007)^18^. Briefly, solvent (99.875% methanol acidified with 1.25% formic acid) was added to fresh tissue material in a 1:3 ratio (fresh weight tissue to solvent), followed by sonication for 15 min at 40 kHz in a water bath at 20°C. The liquid phase was obtained after centrifugation at 15,000 g. Samples were randomly injected (volume: 10 µl) at 20°C into a Vanquish UHPLC with Exploris120 Orbitrap system (Thermo Fisher Scientific, Waltham, MA, USA). The used Vanquish UHPLC was equipped with an Vanquish photodiode array detector (220–600 nm) connected to an Orbitrap Exploris 120 mass spectrometer fitted with an electrospray ionization (ESI) source. A reverse-phase column (Acquity UPLC BEH C18, 1.7 μm, 2.1 × 150 mm; Waters) was used at 40°C for chromatographic separation. Samples were analyzed with a flow rate of 0.4 ml/min with stepwise peak ratios of mobile phase eluents A (0.1% formic acid in ultrapure water), and B (0.1% formic acid in acetonitrile) with initial conditions set to 95:5 (A:B). After the linear gradient of 22 minutes (0.0 – 22.0 min, 5-75% B), washout phases (1min gradient 75% to 90% B, 3 min at 90% B) were followed by recalibration to initial conditions (95%A:5%B). Scans were obtained with an m/z range of 90-1350 Da in positive and negative mode (switched polarity) and a resolution of 60,000 FWMH. Data was obtained in DDA mode (centroid, Number of Dependent Scans = 4) with normalized HCD Collision Energies (%) = 20,40,100. Mass exclusion lists were generated from solvent blank samples injected prior to the tissue sample extracts, to avoid fragmentation of mass features present in the solvent under DDA mode. Quality Control (QC) samples were prepared from pooled samples, including tissue material from all analyzed samples.

### Data processing and downstream analysis

#### MS raw data processing and metabolite feature extraction

Raw Waters MS data was converted to mzXML file format using MSConvert v3.0. Extraction of metabolite features was conducted in mzmine v4.8.0^19^ for a Retention Time (RT) window between 0.8 and 22.8 min. Presets are provided as mzbatch file in *xml* format (see “Data availability section”). Aligned feature lists generated in mzmine v4.8.0 were exported in *mgf* spectra format, and feature quantification table in *csv* format. Principal Component Analysis (PCA) was conducted using *sklearn* v1.7.2 in Python v3.12.0, using *uv* scaling (“autoscaling”,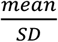). Feature lists generated for spectral data of 14 species, and *mzXML* files of the combined dataset including spectral data of 17 species were submitted to the Global Natural Products Social Molecular Networking online platform using the Feature-Based Molecular Networking (FBMN) and Classical Molecular Networking (CMN) workflows, respectively (GNPS2 Release 2025.12.10)^20^.

#### Structure and compound class annotation

Extracted features of 14 species were annotated in MS2Query v1.0 (De Jonge et al. 2023), DreaMS v1.0^21^, and in SIRIUS v6.1.0^22^, including assessment of prediction accuracy in ZODIAC^23^, and structure annotation by CSI:FingerID^24^. Spectral library searches using DreaMS v1.0 utilized the MassSpecGym spectral library^21^. Compound class annotations based on NP.Classifier^25^ were obtained in CANOPUS^26^ for fingerprints computed in SIRIUS v6.1.0, and for analog predictions in MS2Query v1.0^27^. NP.Classifier annotations were computed via the API (v1.5) for all structure matches and analog predictions obtained with DreaMS v1.0 regardless of similarity scores. Due to the present restriction of the DreaMS model to positive ionization mode training data, DreaMS annotations in negative ionization mode were less reliable. Therefore, only partial matches across MS2Query and SIRIUS were taken into consideration for negative ionization mode compound class annotations. Consensus structure and chemical ontology predictions were computed separately for exact matches of InChI key and compound class strings obtained in SIRIUS v6.1.0 and CANOPUS, and converted from SMILES obtained in DreaMS v1.0 and MS2Query v1.0, using an in-house script available on GitHub See ‘Code availability’ section). Full matches (consensus annotation across all three annotation tools) and partial matches (consensus annotation across at least two out of three tools) per compound class on the NP.Classifier “pathway” level are shown in Fig. 3a. Consensus annotations on the NP.Classifier “superclass” chemical ontology level were obtained for metabolite features in all annotated pathways (Fig.3b).

**Fig. 1.**
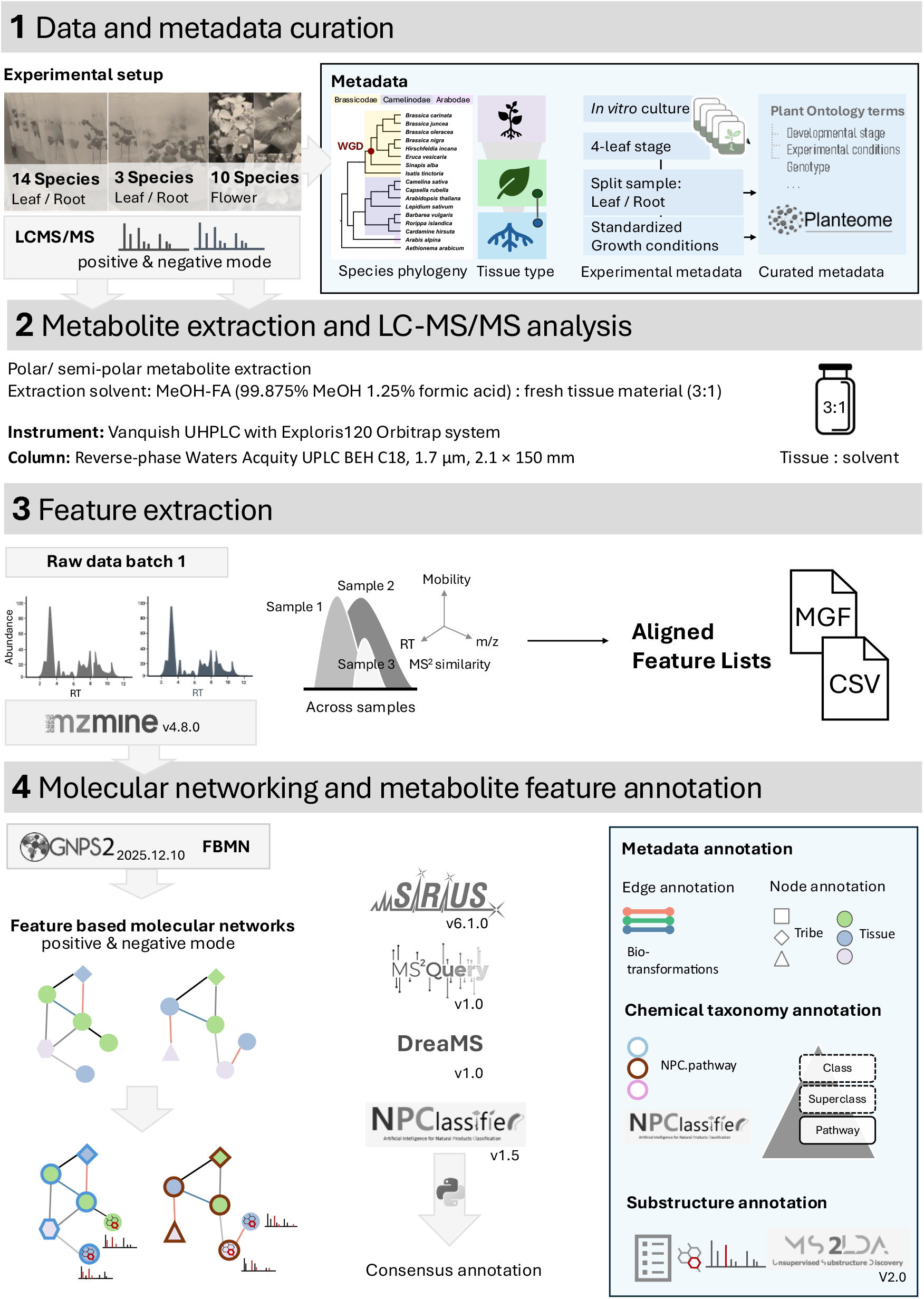
Schematic study design and overview of data acquisition and processing workflow. Plant species and included tissue samples as listed in Table 1. Leaf and root material were obtained from the same sample (split sample). Flower tissue material was harvested from plants grown in the greenhouse. For each sample, tissue material from three nested biological replicates with 5 to 20 plants per replicate were pooled. Feature-Based Molecular Networks (FBMNs), and Classical Molecular Networks (CMNs) were annotated with consensus annotations on the NP.Classifier chemical ontology levels: pathway, superclass, class, along with metadata for species taxonomy, and tissue types.

**Fig. 2.**
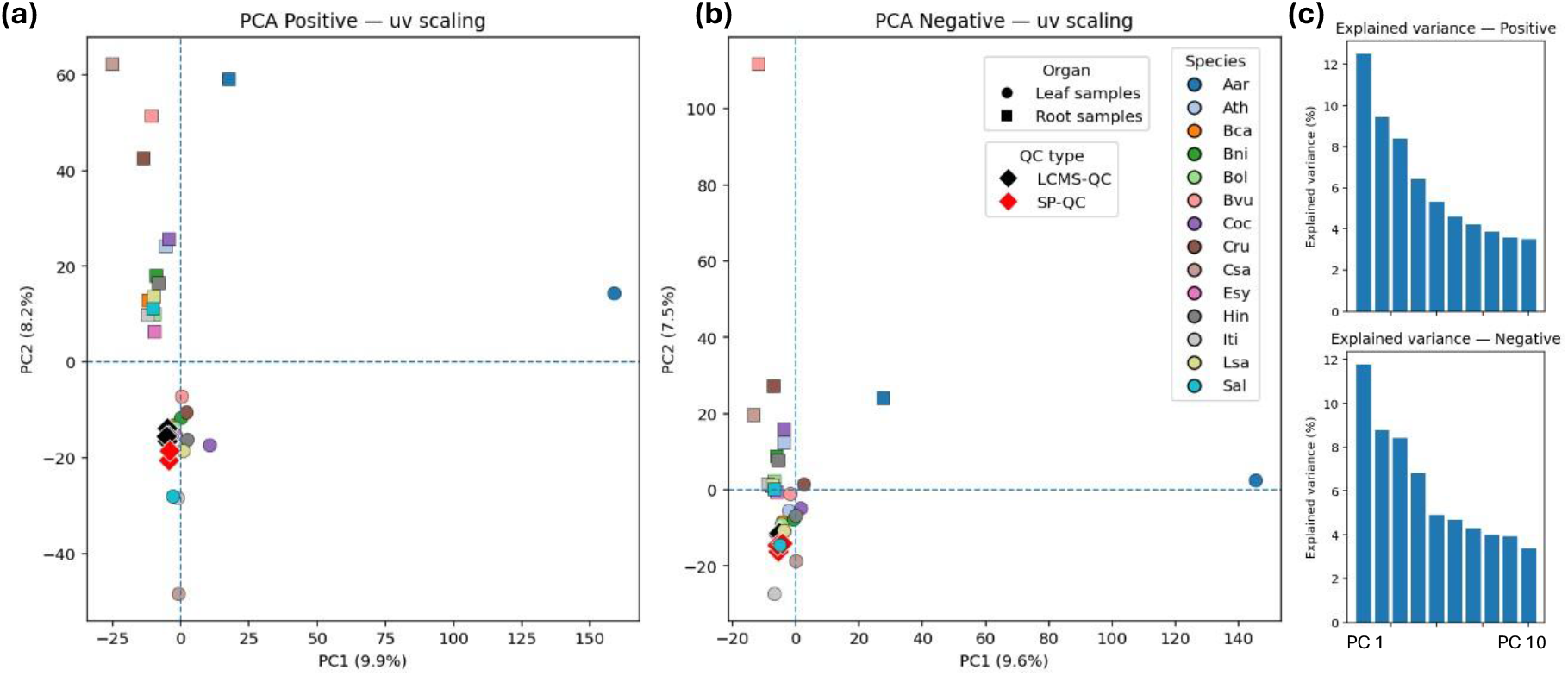
Principal component analysis (PCA) of untargeted LC-MS/MS data in **(a)** positive and **(b)** negative ionization mode. PCA was computed using *uv* scaling (“autoscaling”, 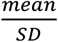) for unfiltered aligned features across 14 species and two tissues (leaf and root), including Quality Control (QC) samples. LCMS-QC samples (n=6) were prepared from extracts of all included samples. SP-QCs (n=2 in positive mode, n=3 in negative mode) were prepared from a pool of fresh tissue material of all included samples, followed by separate extraction. **(c)** Variance explained by principal components (PC1 to PC10).

**Fig. 3.**
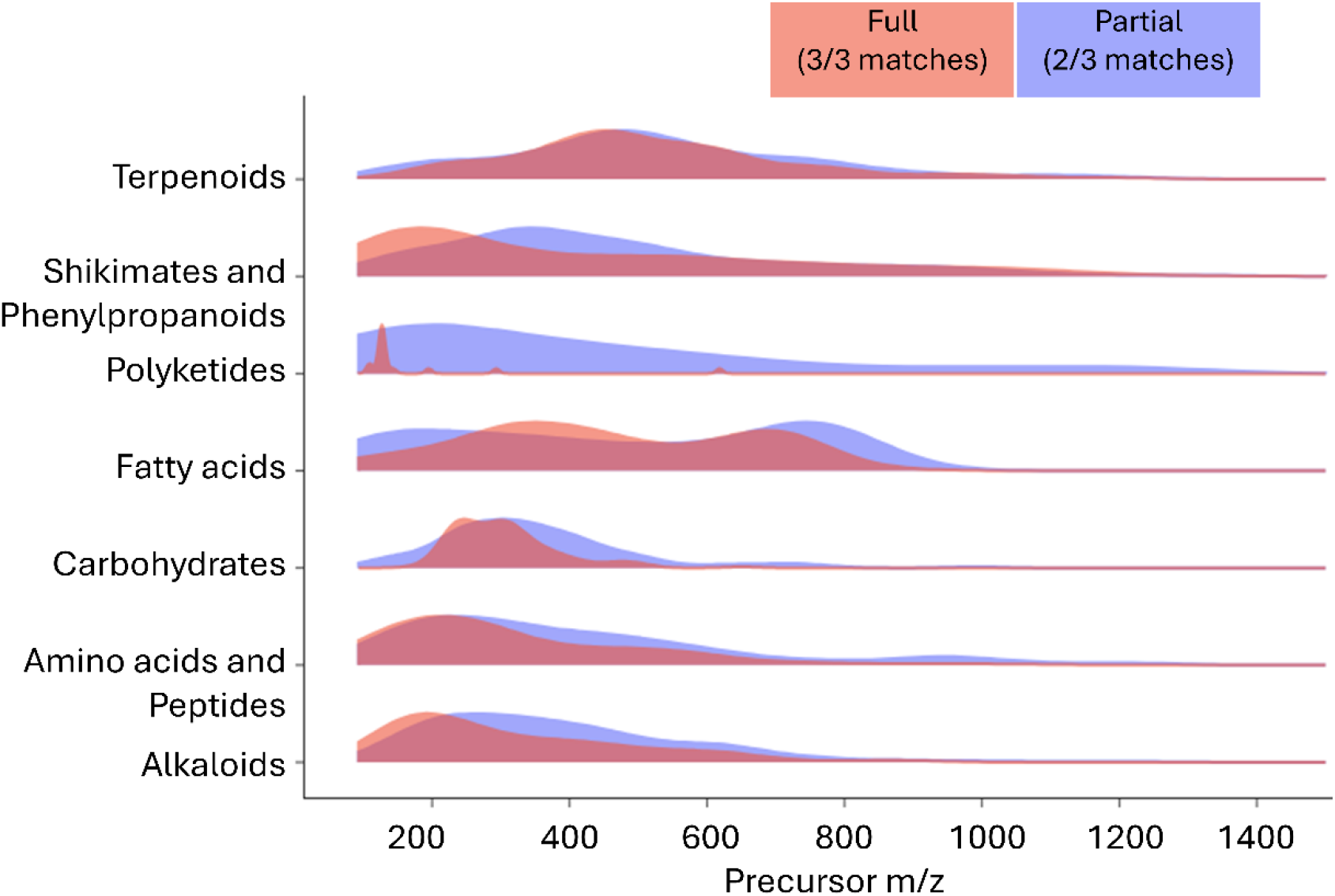
Consensus annotations for NP.Classifier compound classes in positive ionization mode. Full and partial consensus annotations per compound class for precursor m/z values in a range of 120 – 1350 Da.

Total number, compound class categories, and overlap of annotations per annotation tool are available on Zenodo (DOI: 10.5281/zenodo.19391098). Combined output of metabolite feature annotations and consensus predictions was mapped on the FBMN (Fig. 4). The consensus spectra of the CMN obtained for the combined dataset including 17 species was annotated with SIRIUS v6.1.0, using a cutoff of 0.6 for the exact confidence score in downstream analysis.

**Fig. 4.**
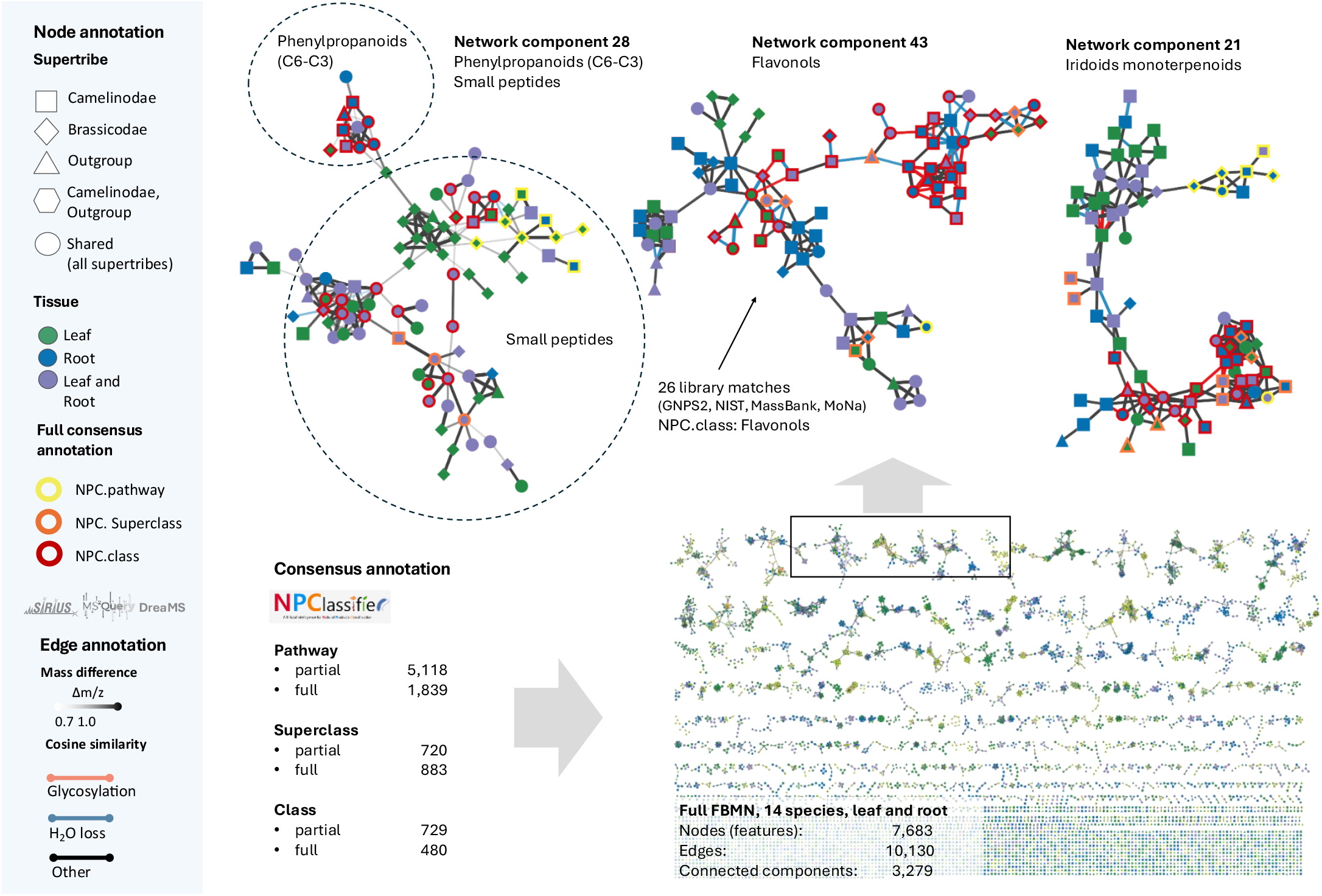
Annotated Feature-Based Molecular Network (FBMN) in positive ionization mode. FBMN of extracted features using mzmine v4.8.0 from untargeted LC-MS/MS data. Sample metadata and consensus annotation on NP.Classifier chemical ontology levels mapped on the FBMN.

### Data Records

#### LC-MS/MS data

Raw scans of MS data for 17 Brassicaceous species (leaf and root tissue) have been deposited in Waters (.*RAW*) and converted *mzXML* format on MetaboLights (MTBLS14270), and corresponding metadata available on Zenodo (DOI: 10.5281/zenodo.19391098), and associated with individual data records has been formatted according to ReDu^28^ (MassIVE ID: MSV000101479) framework standards. Raw data was recorded in positive and negative ionization modes for all samples.

The metabolomics dataset consists of 17 raw spectra datafiles for leaf and root tissue and per ionization mode, respectively. In addition, the dataset includes raw and pre-processed LC-MS/MS data of flower tissue for 10 species in positive and negative mode. In total, 88 sample raw data files are provided for both ionization modes and including all tissues. For quality assessment, 12 QC samples, and two solvent blank injections per batch per ionization mode across two independent LC-MS batches are included. In summary, the dataset consists of 124 raw data files listed in Table 3.

**Table 3.**
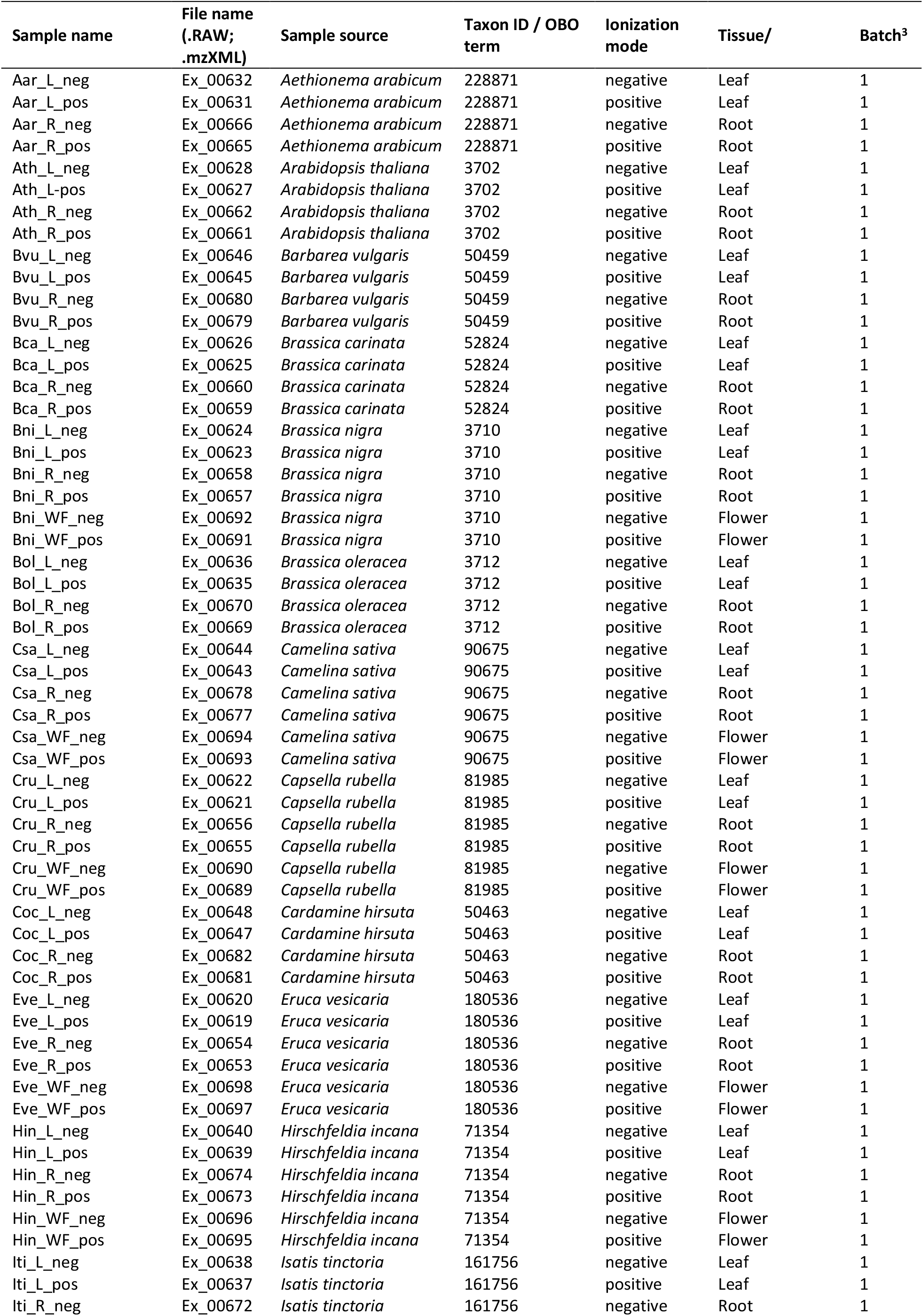
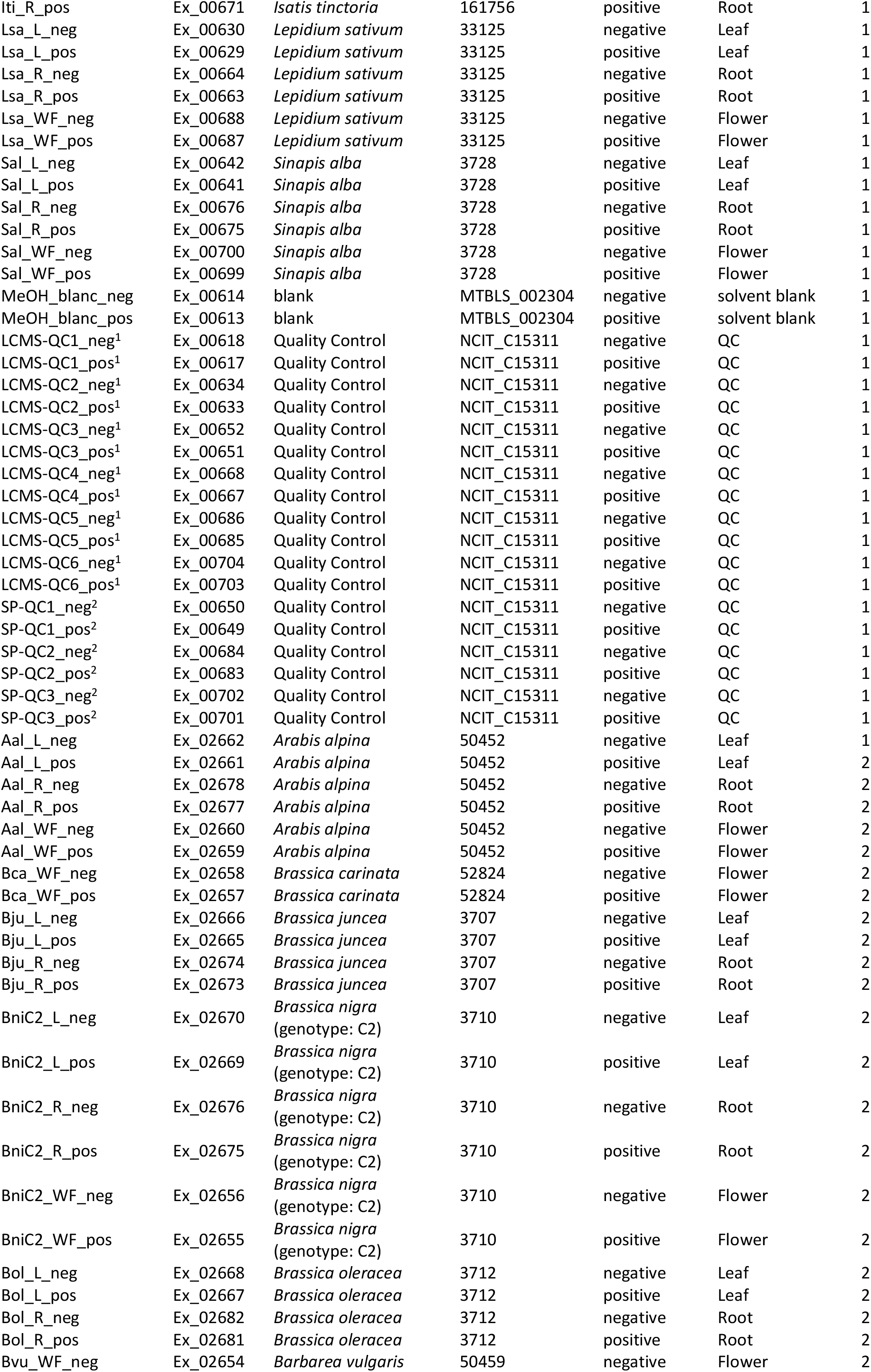
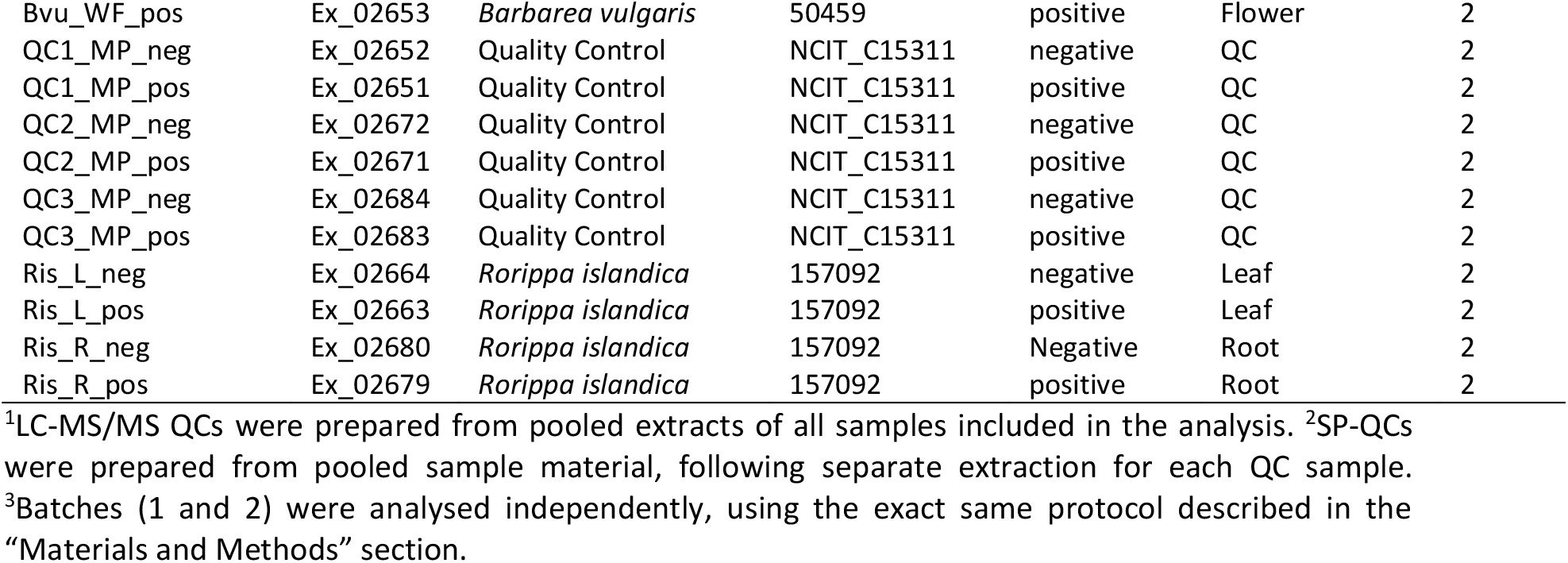
List of untargeted LC-MS/MS data collection across 17 species and three tissues, including Quality Control (QC) samples, and solvent blank samples.

| Sample name | File name<br>(.RAW;<br>.mzXML) | Sample source | Taxon ID / OBO<br>term | Ionization<br>mode | Tissue/ | Batch<sup>3</sup> |
| --- | --- | --- | --- | --- | --- | --- |
| Aar_L_neg | Ex_00632 | *Aethionema arabicum* | 228871 | negative | Leaf | 1 |
| Aar_L_pos | Ex_00631 | *Aethionema arabicum* | 228871 | positive | Leaf | 1 |
| Aar_R_neg | Ex_00666 | *Aethionema arabicum* | 228871 | negative | Root | 1 |
| Aar_R_pos | Ex_00665 | *Aethionema arabicum* | 228871 | positive | Root | 1 |
| Ath_L_neg | Ex_00628 | *Arabidopsis thaliana* | 3702 | negative | Leaf | 1 |
| Ath_L_pos | Ex_00627 | *Arabidopsis thaliana* | 3702 | positive | Leaf | 1 |
| Ath_R_neg | Ex_00662 | *Arabidopsis thaliana* | 3702 | negative | Root | 1 |
| Ath_R_pos | Ex_00661 | *Arabidopsis thaliana* | 3702 | positive | Root | 1 |
| Bvu_L_neg | Ex_00646 | *Barbarea vulgaris* | 50459 | negative | Leaf | 1 |
| Bvu_L_pos | Ex_00645 | *Barbarea vulgaris* | 50459 | positive | Leaf | 1 |
| Bvu_R_neg | Ex_00680 | *Barbarea vulgaris* | 50459 | negative | Root | 1 |
| Bvu_R_pos | Ex_00679 | *Barbarea vulgaris* | 50459 | positive | Root | 1 |
| Bca_L_neg | Ex_00626 | *Brassica carinata* | 52824 | negative | Leaf | 1 |
| Bca_L_pos | Ex_00625 | *Brassica carinata* | 52824 | positive | Leaf | 1 |
| Bca_R_neg | Ex_00660 | *Brassica carinata* | 52824 | negative | Root | 1 |
| Bca_R_pos | Ex_00659 | *Brassica carinata* | 52824 | positive | Root | 1 |
| Bni_L_neg | Ex_00624 | *Brassica nigra* | 3710 | negative | Leaf | 1 |
| Bni_L_pos | Ex_00623 | *Brassica nigra* | 3710 | positive | Leaf | 1 |
| Bni_R_neg | Ex_00658 | *Brassica nigra* | 3710 | negative | Root | 1 |
| Bni_R_pos | Ex_00657 | *Brassica nigra* | 3710 | positive | Root | 1 |
| Bni_WF_neg | Ex_00692 | *Brassica nigra* | 3710 | negative | Flower | 1 |
| Bni_WF_pos | Ex_00691 | *Brassica nigra* | 3710 | positive | Flower | 1 |
| Bol_L_neg | Ex_00636 | *Brassica oleracea* | 3712 | negative | Leaf | 1 |
| Bol_L_pos | Ex_00635 | *Brassica oleracea* | 3712 | positive | Leaf | 1 |
| Bol_R_neg | Ex_00670 | *Brassica oleracea* | 3712 | negative | Root | 1 |
| Bol_R_pos | Ex_00669 | *Brassica oleracea* | 3712 | positive | Root | 1 |
| Csa_L_neg | Ex_00644 | *Camelina sativa* | 90675 | negative | Leaf | 1 |
| Csa_L_pos | Ex_00643 | *Camelina sativa* | 90675 | positive | Leaf | 1 |
| Csa_R_neg | Ex_00678 | *Camelina sativa* | 90675 | negative | Root | 1 |
| Csa_R_pos | Ex_00677 | *Camelina sativa* | 90675 | positive | Root | 1 |
| Csa_WF_neg | Ex_00694 | *Camelina sativa* | 90675 | negative | Flower | 1 |
| Csa_WF_pos | Ex_00693 | *Camelina sativa* | 90675 | positive | Flower | 1 |
| Cru_L_neg | Ex_00622 | *Capsella rubella* | 81985 | negative | Leaf | 1 |
| Cru_L_pos | Ex_00621 | *Capsella rubella* | 81985 | positive | Leaf | 1 |
| Cru_R_neg | Ex_00656 | *Capsella rubella* | 81985 | negative | Root | 1 |
| Cru_R_pos | Ex_00655 | *Capsella rubella* | 81985 | positive | Root | 1 |
| Cru_WF_neg | Ex_00690 | *Capsella rubella* | 81985 | negative | Flower | 1 |
| Cru_WF_pos | Ex_00689 | *Capsella rubella* | 81985 | positive | Flower | 1 |
| Coc_L_neg | Ex_00648 | *Cardamine hirsuta* | 50463 | negative | Leaf | 1 |
| Coc_L_pos | Ex_00647 | *Cardamine hirsuta* | 50463 | positive | Leaf | 1 |
| Coc_R_neg | Ex_00682 | *Cardamine hirsuta* | 50463 | negative | Root | 1 |
| Coc_R_pos | Ex_00681 | *Cardamine hirsuta* | 50463 | positive | Root | 1 |
| Eve_L_neg | Ex_00620 | *Eruca vesicaria* | 180536 | negative | Leaf | 1 |
| Eve_L_pos | Ex_00619 | *Eruca vesicaria* | 180536 | positive | Leaf | 1 |
| Eve_R_neg | Ex_00654 | *Eruca vesicaria* | 180536 | negative | Root | 1 |
| Eve_R_pos | Ex_00653 | *Eruca vesicaria* | 180536 | positive | Root | 1 |
| Eve_WF_neg | Ex_00698 | *Eruca vesicaria* | 180536 | negative | Flower | 1 |
| Eve_WF_pos | Ex_00697 | *Eruca vesicaria* | 180536 | positive | Flower | 1 |
| Hin_L_neg | Ex_00640 | *Hirschfeldia incana* | 71354 | negative | Leaf | 1 |
| Hin_L_pos | Ex_00639 | *Hirschfeldia incana* | 71354 | positive | Leaf | 1 |
| Hin_R_neg | Ex_00674 | *Hirschfeldia incana* | 71354 | negative | Root | 1 |
| Hin_R_pos | Ex_00673 | *Hirschfeldia incana* | 71354 | positive | Root | 1 |
| Hin_WF_neg | Ex_00696 | *Hirschfeldia incana* | 71354 | negative | Flower | 1 |
| Hin_WF_pos | Ex_00695 | *Hirschfeldia incana* | 71354 | positive | Flower | 1 |
| Iti_L_neg | Ex_00638 | *Isatis tinctoria* | 161756 | negative | Leaf | 1 |
| Iti_L_pos | Ex_00637 | *Isatis tinctoria* | 161756 | positive | Leaf | 1 |
| Iti_R_neg | Ex_00672 | *Isatis tinctoria* | 161756 | negative | Root | 1 |
| Iti_R_pos | Ex_00671 | *Isatis tinctoria* | 161756 | positive | Root | 1 |
| Lsa_L_neg | Ex_00630 | *Lepidium sativum* | 33125 | negative | Leaf | 1 |
| Lsa_L_pos | Ex_00629 | *Lepidium sativum* | 33125 | positive | Leaf | 1 |
| Lsa_R_neg | Ex_00664 | *Lepidium sativum* | 33125 | negative | Root | 1 |
| Lsa_R_pos | Ex_00663 | *Lepidium sativum* | 33125 | positive | Root | 1 |
| Lsa_WF_neg | Ex_00688 | *Lepidium sativum* | 33125 | negative | Flower | 1 |
| Lsa_WF_pos | Ex_00687 | *Lepidium sativum* | 33125 | positive | Flower | 1 |
| Sal_L_neg | Ex_00642 | *Sinapis alba* | 3728 | negative | Leaf | 1 |
| Sal_L_pos | Ex_00641 | *Sinapis alba* | 3728 | positive | Leaf | 1 |
| Sal_R_neg | Ex_00676 | *Sinapis alba* | 3728 | negative | Root | 1 |
| Sal_R_pos | Ex_00675 | *Sinapis alba* | 3728 | positive | Root | 1 |
| Sal_WF_neg | Ex_00700 | *Sinapis alba* | 3728 | negative | Flower | 1 |
| Sal_WF_pos | Ex_00699 | *Sinapis alba* | 3728 | positive | Flower | 1 |
| MeOH_blanc_neg | Ex_00614 | blank | MTBLS_002304 | negative | solvent blank | 1 |
| MeOH_blanc_pos | Ex_00613 | blank | MTBLS_002304 | positive | solvent blank | 1 |
| LCMS-QC1_neg<sup>1</sup> | Ex_00618 | Quality Control | NCIT_C15311 | negative | QC | 1 |
| LCMS-QC1_pos<sup>1</sup> | Ex_00617 | Quality Control | NCIT_C15311 | positive | QC | 1 |
| LCMS-QC2_neg<sup>1</sup> | Ex_00634 | Quality Control | NCIT_C15311 | negative | QC | 1 |
| LCMS-QC2_pos<sup>1</sup> | Ex_00633 | Quality Control | NCIT_C15311 | positive | QC | 1 |
| LCMS-QC3_neg<sup>1</sup> | Ex_00652 | Quality Control | NCIT_C15311 | negative | QC | 1 |
| LCMS-QC3_pos<sup>1</sup> | Ex_00651 | Quality Control | NCIT_C15311 | positive | QC | 1 |
| LCMS-QC4_neg<sup>1</sup> | Ex_00668 | Quality Control | NCIT_C15311 | negative | QC | 1 |
| LCMS-QC4_pos<sup>1</sup> | Ex_00667 | Quality Control | NCIT_C15311 | positive | QC | 1 |
| LCMS-QC5_neg<sup>1</sup> | Ex_00686 | Quality Control | NCIT_C15311 | negative | QC | 1 |
| LCMS-QC5_pos<sup>1</sup> | Ex_00685 | Quality Control | NCIT_C15311 | positive | QC | 1 |
| LCMS-QC6_neg<sup>1</sup> | Ex_00704 | Quality Control | NCIT_C15311 | negative | QC | 1 |
| LCMS-QC6_pos<sup>1</sup> | Ex_00703 | Quality Control | NCIT_C15311 | positive | QC | 1 |
| SP-QC1_neg<sup>2</sup> | Ex_00650 | Quality Control | NCIT_C15311 | negative | QC | 1 |
| SP-QC1_pos<sup>2</sup> | Ex_00649 | Quality Control | NCIT_C15311 | positive | QC | 1 |
| SP-QC2_neg<sup>2</sup> | Ex_00684 | Quality Control | NCIT_C15311 | negative | QC | 1 |
| SP-QC2_pos<sup>2</sup> | Ex_00683 | Quality Control | NCIT_C15311 | positive | QC | 1 |
| SP-QC3_neg<sup>2</sup> | Ex_00702 | Quality Control | NCIT_C15311 | negative | QC | 1 |
| SP-QC3_pos<sup>2</sup> | Ex_00701 | Quality Control | NCIT_C15311 | positive | QC | 1 |
| Aal_L_neg | Ex_02662 | *Arabis alpina* | 50452 | negative | Leaf | 1 |
| Aal_L_pos | Ex_02661 | *Arabis alpina* | 50452 | positive | Leaf | 2 |
| Aal_R_neg | Ex_02678 | *Arabis alpina* | 50452 | negative | Root | 2 |
| Aal_R_pos | Ex_02677 | *Arabis alpina* | 50452 | positive | Root | 2 |
| Aal_WF_neg | Ex_02660 | *Arabis alpina* | 50452 | negative | Flower | 2 |
| Aal_WF_pos | Ex_02659 | *Arabis alpina* | 50452 | positive | Flower | 2 |
| Bca_WF_neg | Ex_02658 | *Brassica carinata* | 52824 | negative | Flower | 2 |
| Bca_WF_pos | Ex_02657 | *Brassica carinata* | 52824 | positive | Flower | 2 |
| Bju_L_neg | Ex_02666 | *Brassica juncea* | 3707 | negative | Leaf | 2 |
| Bju_L_pos | Ex_02665 | *Brassica juncea* | 3707 | positive | Leaf | 2 |
| Bju_R_neg | Ex_02674 | *Brassica juncea* | 3707 | negative | Root | 2 |
| Bju_R_pos | Ex_02673 | *Brassica juncea* | 3707 | positive | Root | 2 |
| BniC2_L_neg | Ex_02670 | *Brassica nigra*<br>(genotype: C2) | 3710 | negative | Leaf | 2 |
| BniC2_L_pos | Ex_02669 | *Brassica nigra*<br>(genotype: C2) | 3710 | positive | Leaf | 2 |
| BniC2_R_neg | Ex_02676 | *Brassica nigra*<br>(genotype: C2) | 3710 | negative | Root | 2 |
| BniC2_R_pos | Ex_02675 | *Brassica nigra*<br>(genotype: C2) | 3710 | positive | Root | 2 |
| BniC2_WF_neg | Ex_02656 | *Brassica nigra*<br>(genotype: C2) | 3710 | negative | Flower | 2 |
| BniC2_WF_pos | Ex_02655 | *Brassica nigra*<br>(genotype: C2) | 3710 | positive | Flower | 2 |
| Bol_L_neg | Ex_02668 | *Brassica oleracea* | 3712 | negative | Leaf | 2 |
| Bol_L_pos | Ex_02667 | *Brassica oleracea* | 3712 | positive | Leaf | 2 |
| Bol_R_neg | Ex_02682 | *Brassica oleracea* | 3712 | negative | Root | 2 |
| Bol_R_pos | Ex_02681 | *Brassica oleracea* | 3712 | positive | Root | 2 |
| Bvu_WF_neg | Ex_02654 | *Barbarea vulgaris* | 50459 | negative | Flower | 2 |
| Bvu_WF_pos | Ex_02653 | *Barbarea vulgaris* | 50459 | positive | Flower | 2 |
| QC1_MP_neg | Ex_02652 | Quality Control | NCIT_C15311 | negative | QC | 2 |
| QC1_MP_pos | Ex_02651 | Quality Control | NCIT_C15311 | positive | QC | 2 |
| QC2_MP_neg | Ex_02672 | Quality Control | NCIT_C15311 | negative | QC | 2 |
| QC2_MP_pos | Ex_02671 | Quality Control | NCIT_C15311 | positive | QC | 2 |
| QC3_MP_neg | Ex_02684 | Quality Control | NCIT_C15311 | negative | QC | 2 |
| QC3_MP_pos | Ex_02683 | Quality Control | NCIT_C15311 | positive | QC | 2 |
| Ris_L_neg | Ex_02664 | *Rorippa islandica* | 157092 | negative | Leaf | 2 |
| Ris_L_pos | Ex_02663 | *Rorippa islandica* | 157092 | positive | Leaf | 2 |
| Ris_R_neg | Ex_02680 | *Rorippa islandica* | 157092 | Negative | Root | 2 |
| Ris_R_pos | Ex_02679 | *Rorippa islandica* | 157092 | positive | Root | 2 |

<sup>1</sup>LC-MS/MS QCs were prepared from pooled extracts of all samples included in the analysis. <sup>2</sup>SP-QCs were prepared from pooled sample material, following separate extraction for each QC sample.
<sup>3</sup>Batches (1 and 2) were analysed independently, using the exact same protocol described in the “Materials and Methods” section.

**Table 4.** Intermediate processing files for untargeted LC-MS/MS analysis in positive and negative ionization mode.

| Description | Filename on Zenodo |
| --- | --- |
| Batchfile for mzmine v4.8.0<br>(negative ionization mode) | batch_MP23_neg_4-8-0_v4_01a_251024.mzbatch |
| Batchfile for mzmine v4.8.0<br>(positive ionization mode) | batch_MP23_pos_4-8-0_v4_01a_251024.mzbatch |
| Aligned feature lists for 14 species, leaf and root tissue (negative ionization mode) | 14_LR_neg_4-8-0_v4_01a_251024_Aligned_feature_list.mgf |
| Quantification table for aligned feature list (negative ionization mode) | 14_LR_neg_4-8-0_v4_01a_251024_Aligned_feature_list_quant.csv |
| Aligned feature lists for 14 species, leaf and root tissue (positive ionization mode) | 14LR_pos_mzbatch_v3_4-8-0_251117_Aligned_feature_list |
| Quantification table for aligned feature list (positive ionization mode) | 14LR_pos_mzbatch_v3_4-8-0_251117_Aligned_feature_list_quant.csv |
| Complete results for MS2LDA (negative ionization mode) | ms2lda_14LR_neg_260114 (folder) |
| Complete results for MS2LDA (positive ionization mode) | ms2lda_14LR_pos_260113 (folder) |
| Fully annotated FBMNs in Cytoscape (session file) | 14_LR_pos_neg_Full_networks_annotated.cys |
| Consensus MGF file output of MSCluster, including 17 species (leaf and root) (negative ionization mode) | 17LR_neg_consensus-spectra_MSCluster_specs_ms_GNPS2.mgf |
| Complete summary of SIRIUS v6.1.0 annotations for consensus MGF including 17 species (leaf and root) in (negative ionization mode) | 2507101_SIRIUS_17_LR_neg_250710_SIRIUS_Summary (folder) |

#### Annotation tables and mappings

Aligned feature lists and feature quantification tables computed in mzmine v4.8.0, IDs of GNPS2 workflows including all processing parameters, associated metadata, extracted Mass2Motifs, along with fully annotated Feature-Based Molecular Networks (FBMNs) including all annotations were deposited on Zenodo (DOI: 10.5281/zenodo.19391098).

### Technical validation

For calibration and stabilisation of retention time in the LC-system, the first sample was injected four times at the beginning of the serial analysis. Mass exclusion lists for fragmentation during the sample series analysis were generated based on analysis of a blank solvent injection to prevent fragmentation of mass features abundant in solvent blanks. Variation, degradation and adduct formation over time were assessed using three Quality Control (QC) samples, consisting of independently prepared pooled sample material (SP-QC), and three QC samples pooled from all prepared extracts (LCMS-QC). SP-QCs were used to account for variation in sample preparation and extraction, and LCMS-QCs were used to separately assess variation in analytical measurements. One SP-QC replicate in positive ionization mode (Sp-QC1_pos_Ex_00649.mzXML) has been excluded from technical validation due to low quality of the chromatogram. Across LCMS-QCs and SP-QCs, 1118 (63.5%) and 814 (59.8%) of all detected metabolite features overlapped in positive and negative ionization mode, respectively (Fig. 5a), while total numbers of detected features were generally lower in SP-QC samples (Fig. 5b).

**Fig. 5.**
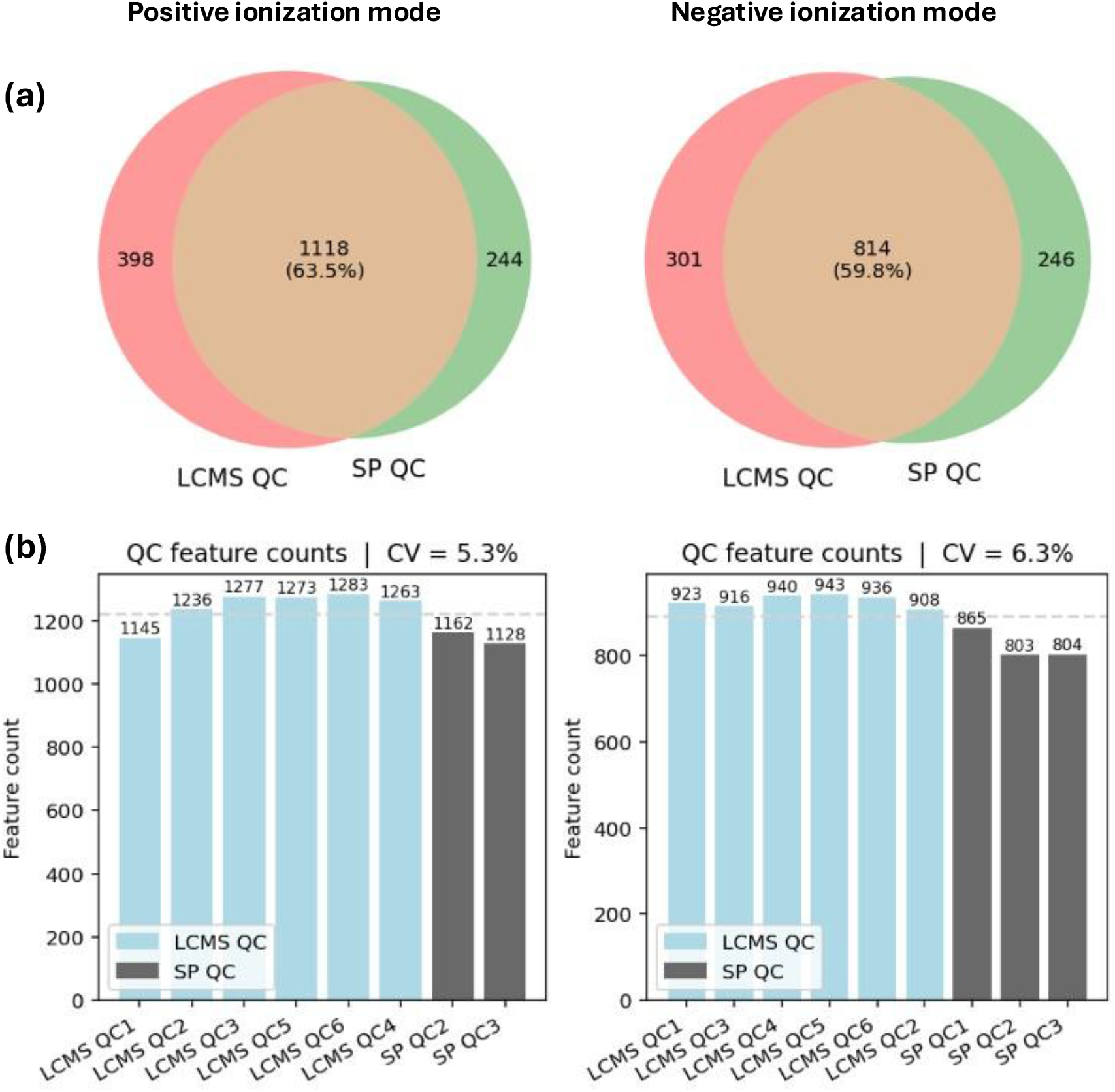
Overlap of detected features, and total feature counts in QC sampled in positive and negative ionization mode. **(a)** Venn diagram showing overlap of features between SP-QC and LCMS-QC samples. **(b)** Total number of features extracted from each individual QC sample. SP-QC samples were prepared individually, including full extraction from pooled sample (fresh tissue) material. In positive ionization mode SP-QC1 has been excluded from the analysis. LCMS-QC samples were prepared from pooled sample extracts.

In the PCA using Pareto scaling 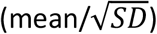, LCMS-QCs and SP-QCs were mostly separated from leaf and root tissue samples under positive ionization mode, with leaf samples clustering more closely together than root samples (Fig. 6a). Clear separation of LCMS-QCs and SP-QCs can be observed only in negative ionization mode (Fig. 6b), while loadings (individual mass features) did not form separate clusters in either ionization mode.

**Fig. 6.**
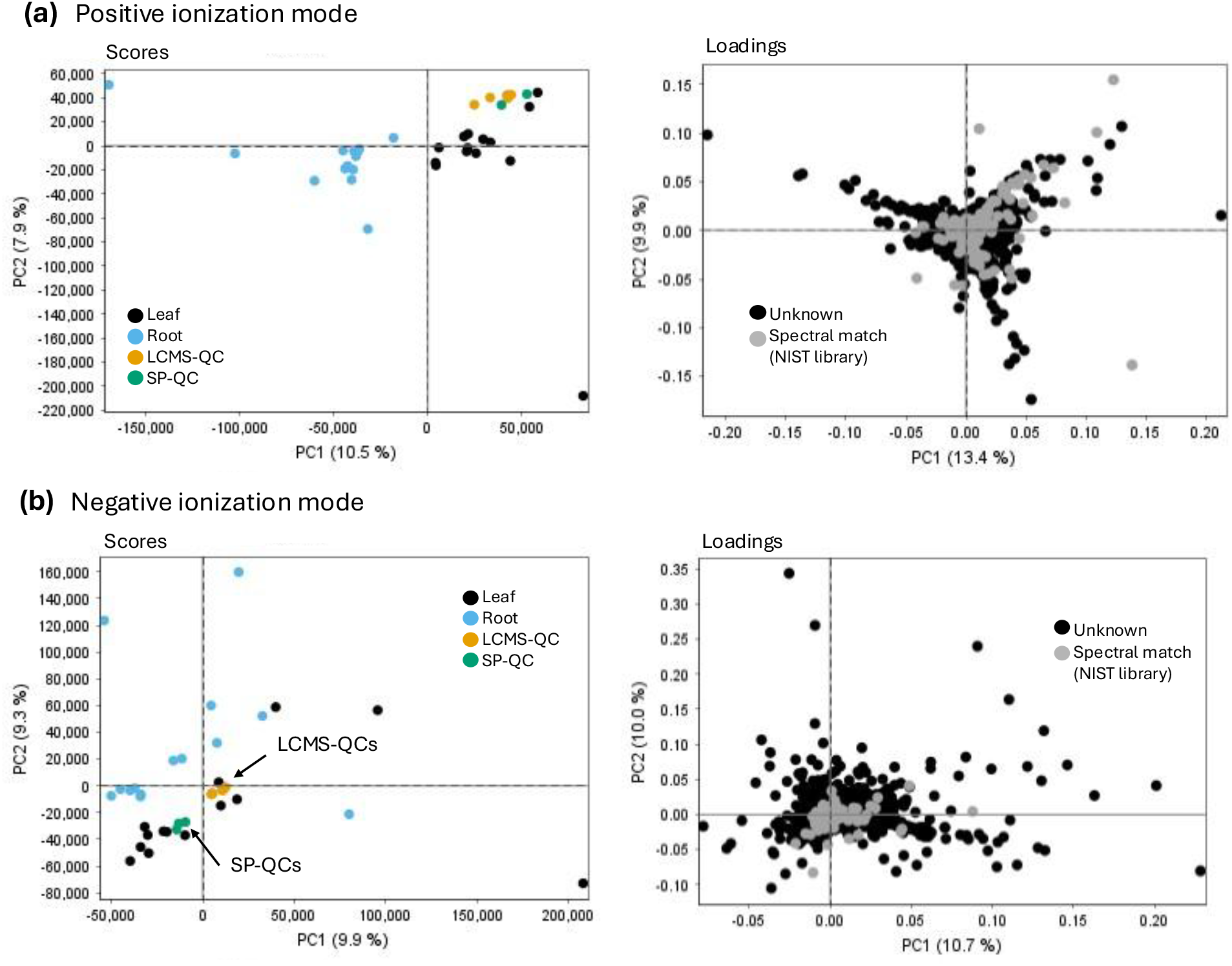
PCA scores and loadings of aligned feature lists of 14 species, including LCMS-QCs (pooled from sample extracts) and SP-QCs (pooled from fresh tissue material of all samples). **(a)** Features extracted in positive ionization mode and **(b)** in negative ionization mode. Samples were coloured based on source tissue material (leaf, root).

Overlap of detected features in LCMS-QCs and SP-QCs with all included samples was found mostly unbiased for included species and tissues, indicating that samples with higher total numbers of detected features were not overrepresented in the QC samples (Fig. 7). In positive ionization mode, 1,034 features overlapped with both QC groups and across all samples, while the overlap was slightly higher in LCMS-QCs (1,333 features) compared to SP-QCs (1,218 features, Fig. 7a). Similarly, in negative ionization mode, 736 mass features overlapped between both QC groups and samples, while 963 mass features were shared between samples and LCMS-QCs, and 897 mass features were shared between samples and SP-QCs (Fig 7b). Across both QC groups, overlap of all detected features in a sample and the QCs ranges from 27.93% to 67.35%, with higher mean overlap in LCMS-QCs (48.62%) compared to SP-QCs (45.42%).

**Fig. 7.**
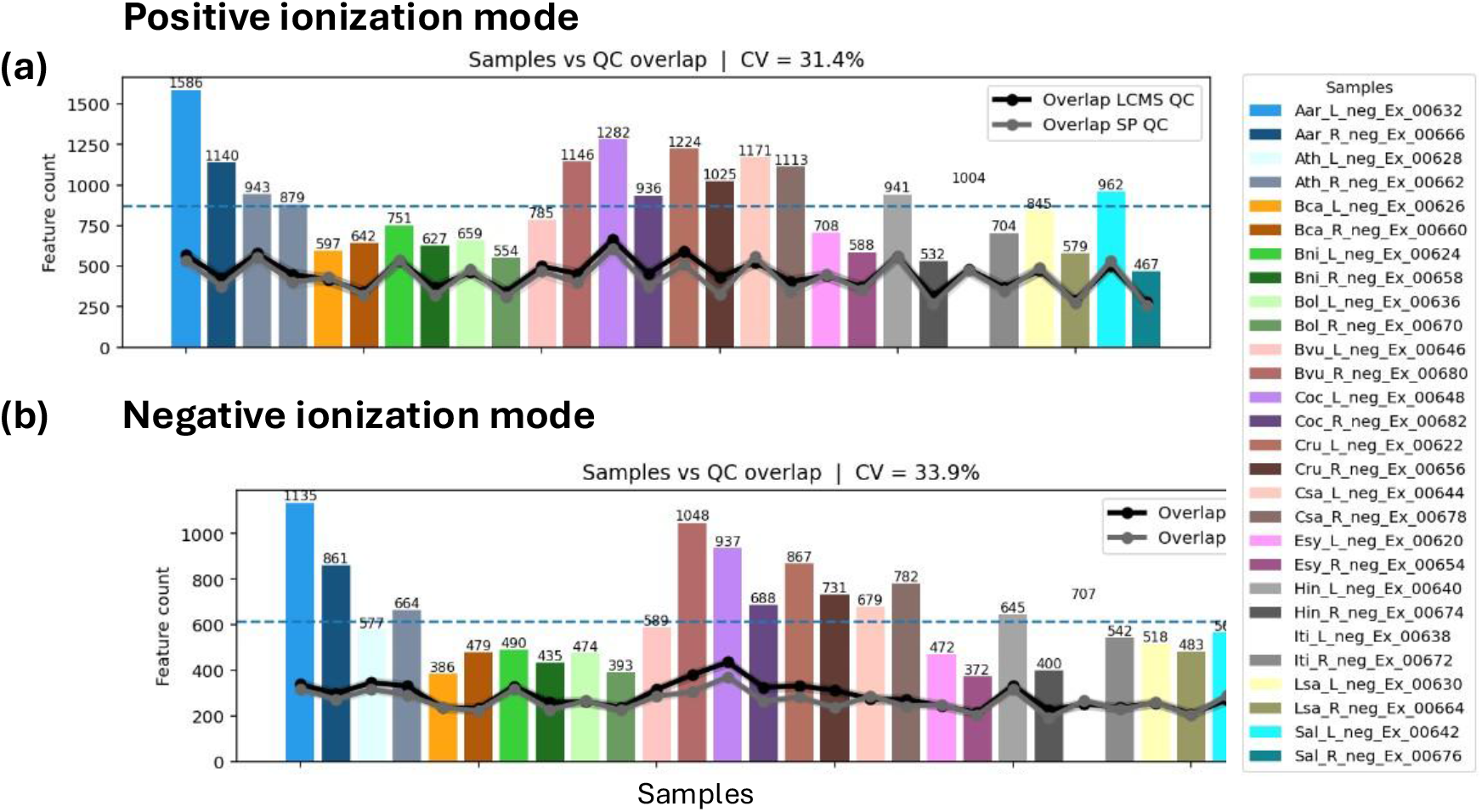
Overlaps of QC samples with each individual sample included in untargeted LC-MS/MS analysis in **(a)** positive and **(b)** negative ionization mode. Total numbers of features detected in each sample, and overlap of detected features in SP-QC and LCMS-QC samples (mean) following feature extraction and alignment. Blue dashed line shows mean overlap.

### Usage notes

The presented dataset provides untargeted metabolomics data suitable for correlation-based omics data integration, and genome assemblies are available for all included species on public repositories (NCBI, JGI, Cruciferseq), as listed in Table 1 (Methods). Raw data processing workflows and intermediate output files are provided for each processing step, to facilitate downstream processing with alternative software, and version control following software updates. Processing workflows for untargeted LC-MS/MS data have been designed with consideration to recent developments of established software in the respective field^29-30^. Comparison of intermediate file processing and data transformation for workflow modules is facilitated by supplementary output summaries provided on Zenodo (DOI: 10.5281/zenodo.19391098) The documentation of MZmine v4.8.5 batchfiles is provided for each processing module to aid adaptations of individual batch processing steps.

## Code availability

Code for transformation of feature list annotations, visualization, annotation and manipulation of phylogenetic trees are provided in R v4.3.2. Scripts for computation of consensus compound class annotations are written in Python v3.12. All custom scripts are provided on GitHub (github.com/capsicumbaccatum/Manuscript_Metabolic_profiling). Intermediate outputs and configuration files (exported feature lists, mzmine batchfiles) are available on Zenodo (DOI: 10.5281/zenodo.19391098). Custom code written in Python v3.12.0 employed the following packages: RDKit v2025_09_3^31^ for SMILES string conversion and Matchms^32^ and MS2LDA 2.0^33-35^ for visualization of mass spectra and for motif extraction and formatting.

## Acknowledgements

This work was supported by the Netherlands Organization for Scientific Research (NWO) Vidi Grant VI.Vidi.213.183 to M.H.M. The authors would like to thank Ric de Vos and Bert Schipper for their help and comments on the experimental design and metabolite extraction protocols, and for establishing the analytical platforms.

## Author contributions

F.W., M.H.M., K.B., and J.J.J.vdH. designed the study. F.W. and T.W. conducted the experiments and collected the data. F.W. and J.J.J.vdH. conducted LC-MS/MS data analysis. F.W. created figures and tables. F.W. wrote the initial draft of the manuscript, and all authors contributed substantially to revisions. No generative AI (LLMs) were used for writing this manuscript.

## References

1. Singh, K. S., van der Hooft, J. J. J., van Wees, S. C. M. & Medema, M. H. Integrative omics approaches for biosynthetic pathway discovery in plants. Nat. Prod. Rep. 39, 1876–1896 (2022).

2. Alseekh, S. & Fernie, A. R. Expanding our coverage: Strategies to detect a greater range of metabolites. Curr. Opin. Plant Biol. 73, 102335 (2023).

3. Alseekh, S. & Fernie, A. R. Metabolomics 20 years on: what have we learned and what hurdles remain? Plant J. 94, 933–942 (2018).

4. Walden, N. et al. Nested whole-genome duplications coincide with diversification and high morphological disparity in Brassicaceae. Nat. Commun. 11, 3795 (2020).

5. Hendriks, K. P., Kiefer, C., Al-Shehbaz, I. A., Bailey, C. D., Hooft van Huysduynen, A., Nikolov, L. A., … & Lens, F. Global phylogeny of the Brassicaceae provides important insights into gene discordance. bioRxiv, 2022–09 (2022).

6. Tohge, T. et al. Functional genomics by integrated analysis of metabolome and transcriptome of Arabidopsis plants over-expressing an MYB transcription factor: Metabolomics and transcriptomics. Plant J. 42, 218–235 (2005).

7. Kang, K. B. et al. Comprehensive mass spectrometry−guided phenotyping of plant specialized metabolites reveals metabolic diversity in the cosmopolitan plant family Rhamnaceae. Plant J. 98, 1134–1144 (2019).

8. Padilla-González, G. F., Rosselli, A., Sadgrove, N. J., Cui, M. & Simmonds, M. S. J. Mining the chemical diversity of the hemp seed (*Cannabis sativa* L.) metabolome: discovery of a new molecular family widely distributed across hemp. Front. Plant Sci. 14, 1114398 (2023).

9. Harun, S., Abdullah-Zawawi, M.-R., Goh, H.-H. & Mohamed-Hussein, Z.-A. A Comprehensive Gene Inventory for Glucosinolate Biosynthetic Pathway in *Arabidopsis thaliana*. J. Agric. Food Chem. 68, 7281–7297 (2020).

10. Agerbirk, N., Hansen, C. C., Kiefer, C., Hauser, T. P., Ørgaard, M., Lange, C. B. A., … & Koch, M. A. Comparison of glucosinolate diversity in the crucifer tribe Cardamineae and the remaining order Brassicales highlights repetitive evolutionary loss and gain of biosynthetic steps. Phytochemistry, 185, 112668 (2021).

11. Bednarek, P., Piślewska−Bednarek, M., Ver Loren van Themaat, E., Maddula, R. K., Svatoš, A., & Schulze−Lefert, P. Conservation and clade−specific diversification of pathogen− inducible tryptophan and indole glucosinolate metabolism in *Arabidopsis thaliana* relatives. New Phytologist, 192(3), 713–726 (2011).

12. German, D. A., Hendriks, K. P., Koch, M. A., Lens, F., Lysak, M. A., Bailey, C. D., … & Al-Shehbaz, I. A. An updated classification of the Brassicaceae (*Cruciferae*). PhytoKeys, 220, 127 (2023).

13. Barrett, T., Wilhite, S. E., Ledoux, P., Evangelista, C., Kim, I. F., Tomashevsky, M., Marshall, K. A., Phillippy, K. H., Sherman, P. M., Holko, M., Yefanov, A., Lee, H., Zhang, N., Robertson, C. L., Serova, N., Davis, S., & Soboleva, A. NCBI GEO: archive for functional genomics data sets--update. Nucleic acids research, 41 (Database issue), D991–D995 (2013).

14. Sayers, E. W., Beck, J., Bolton, E. E., Brister, J. R., Chan, J., Connor, R., … & Pruitt, K. D. Database resources of the National Center for Biotechnology Information in 2025. Nucleic acids research, 53(D1), D20–D29 (2025).

15. Allard, P. M., Gaudry, A., Quirós-Guerrero, L. M., Rutz, A., Dounoue-Kubo, M., Walker, T. W., … & Wolfender, J. L. Open and reusable annotated mass spectrometry dataset of a chemodiverse collection of 1,600 plant extracts. GigaScience, 12, giac124 (2023).

16. Yurekten, O., Payne, T., Tejera, N., Amaladoss, F. X., Martin, C., Williams, M., & O’Donovan, C. MetaboLights: open data repository for metabolomics. Nucleic acids research, 52(D1), D640–D646 (2024).

17. El Abiead, Y., Strobel, M., Payne, T., Fahy, E., O’Donovan, C., Subramamiam, S., … & Wang, M. Enabling pan-repository reanalysis for big data science of public metabolomics data. Nature Communications, 16(1), 4838 (2025).

18. De Vos, R. C. et al. Untargeted large-scale plant metabolomics using liquid chromatography coupled to mass spectrometry. Nat. Protoc. 2, 778–791 (2007).

19. Schmid, R., Heuckeroth, S., Korf, A., Smirnov, A., Myers, O., Dyrlund, T. S., … & Pluskal, T. Integrative analysis of multimodal mass spectrometry data in MZmine 3. Nature biotechnology, 41(4), 447–449 (2023).

20. Wang, M., Carver, J. J., Phelan, V. V., Sanchez, L. M., Garg, N., Peng, Y., … & Bandeira, N.. Sharing and community curation of mass spectrometry data with Global Natural Products Social Molecular Networking. Nature biotechnology, 34(8), 828–837 (2016).

21. Bushuiev, R., Bushuiev, A., Samusevich, R., Brungs, C., Sivic, J., & Pluskal, T. Self-supervised learning of molecular representations from millions of tandem mass spectra using DreaMS. Nature Biotechnology, 1–11 (2025).

22. Dührkop, K., Nothias, L. F., Fleischauer, M., Reher, R., Ludwig, M., Hoffmann, M. A., … & Böcker, S. Systematic classification of unknown metabolites using high-resolution fragmentation mass spectra. Nature biotechnology, 39(4), 462–471 (2021).

23. Ludwig, M., Nothias, L. F., Dührkop, K., Koester, I., Fleischauer, M., Hoffmann, M. A., … & Böcker, S. Database-independent molecular formula annotation using Gibbs sampling through ZODIAC. Nature Machine Intelligence, 2(10), 629–641 (2020).

24. Dührkop, K., Shen, H., Meusel, M., Rousu, J. & Böcker, S. Searching molecular structure databases with tandem mass spectra using CSI:FingerID. Proc. Natl. Acad. Sci. 112, 12580–12585 (2015).

25. Kim, H. W., Wang, M., Leber, C. A., Nothias, L. F., Reher, R., Kang, K. B., … & Cottrell, G. W. NPClassifier: a deep neural network-based structural classification tool for natural products. Journal of natural products, 84(11), 2795–2807 (2021).

26. Dührkop, K., Nothias, L. F., Fleischauer, M., Reher, R., Ludwig, M., Hoffmann, M. A., … & Böcker, S. Systematic classification of unknown metabolites using high-resolution fragmentation mass spectra. Nature biotechnology, 39(4), 462–471 (2021).

27. de Jonge, N. F., Louwen, J. J., Chekmeneva, E., Camuzeaux, S., Vermeir, F. J., Jansen, R. S., … & van der Hooft, J. J. MS2Query: reliable and scalable MS2 mass spectra-based analogue search. Nature Communications, 14(1), 1752 (2023).

28. Jarmusch, A. K., Wang, M., Aceves, C. M., Advani, R. S., Aguirre, S., Aksenov, A. A., … & Dorrestein, P. C. ReDU: a framework to find and reanalyze public mass spectrometry data. Nature Methods, 17(9), 901–904 (2020).

29. Cai, Y., Zhou, Z. & Zhu, Z.-J. Advanced analytical and informatic strategies for metabolite annotation in untargeted metabolomics. TrAC Trends Anal. Chem. 158, 116903 (2023).

30. Alseekh, S., Aharoni, A., Brotman, Y., Contrepois, K., D’Auria, J., Ewald, J., … & Fernie, A. R. Mass spectrometry-based metabolomics: a guide for annotation, quantification and best reporting practices. Nature methods, 18(7), 747–756 (2021).

31. Landrum, G., Tosco, P., Kelley, B., Rodriguez, R., Cosgrove, D., Vianello, R., … & Monat, J. rdkit/rdkit: 2025_03_1 (Q1 2025) Release. Zenodo (2025).

32. Huber, F., Verhoeven, S., Meijer, C., Spreeuw, H., Castilla, E. M. V., Geng, C., … & Spaaks, J. H. Matchms-processing and similarity evaluation of mass spectrometry data. BioRxiv, 2020–08 (2020).

33. Wandy, J., Zhu, Y., van der Hooft, J. J., Daly, R., Barrett, M. P., & Rogers, S. Ms2lda. org: web-based topic modelling for substructure discovery in mass spectrometry. Bioinformatics, 34(2), 317–318 (2018).

34. van Der Hooft, J. J. J., Wandy, J., Barrett, M. P., Burgess, K. E., & Rogers, S. Topic modeling for untargeted substructure exploration in metabolomics. Proceedings of the National Academy of Sciences, 113(48), 13738–13743 (2016).

35. Ortega, L. R. T., Dietrich, J., Wandy, J., Mol, H., & van der Hooft, J. J. Large-scale discovery and annotation of hidden substructure patterns in mass spectrometry profiles. bioRxiv, 2025–06 (2025).

